# UPS-indel: a Universal Positioning System for Indels

**DOI:** 10.1101/133553

**Authors:** Mohammad Shabbir Hasan, Xiaowei Wu, Layne T. Watson, Zhiyi Li, Liqing Zhang

## Abstract

**Background:** Indels, though differing in allele sequence and position, are biologically equivalent when they lead to the same altered sequences. Storing biologically equivalent indels as distinct entries in databases causes data redundancy, and may mislead downstream analysis and interpretations. About 10% of the human indels stored in dbSNP are redundant. It is thus desirable to have a unified system for identifying and representing equivalent indels in publically available databases. Moreover, a unified system is also desirable to compare the indel calling results produced by different tools. This paper describes UPS-indel, a utility tool that creates a universal positioning system for indels so that equivalent indels can be uniquely determined by their coordinates in the new system, which also can be used to compare indel calling results produced by different tools.

**Results:** UPS-indel identifies nearly 15% indels in dbSNP (version 142) as redundant across all human chromosomes, higher than previously reported. When applied to COSMIC coding and noncoding indel datasets, UPS-indel identifies nearly 29% and 13% indels as redundant, respectively. Comparing the performance of UPS-indel with existing variant normalization tools vt normalize, BCFtools, and GATK LeftAlignAndTrimVariants shows that UPS-indel is able to identify 456,352 more redundant indels in dbSNP; 2,118 more in COSMIC coding, and 553 more in COSMIC noncoding indel dataset in addition to the ones reported jointly by these tools. Moreover, comparing UPS-indel to other state-of-the-art approaches for indel call set comparison demonstrates that UPS-indel is clearly superior to other approaches in finding indels in common among call sets.

**Conclusions:** UPS-indel is theoretically proven to find all equivalent indels, and is thus exhaustive. UPS-indel is written in C++ and the command line version is freely available to download at http://ups-indel.sourceforge.net. The online version of UPS-indel is available at http://bench.cs.vt.edu/ups-indel/.

## Background

Indel stands for insertion or deletion of bases in a DNA sequence. As the second most common form of genetic variation, indels play an important role in genome and protein evolution. Due to artificial factors such as sequencing errors, ambiguous alignment of the reads, inconsistent ways of representing the same variant by different tools, the same mutation may be recognized as distinct variations occurring at different locations [1–3]. For example, consider a reference sequence AGGAAAGAAAGAAAGAAAGAG ranging from position 100285630 to 100285650 and two indels stored in dbSNP, rs147659011 (GAAA/+) and rs60376183 (AAGA/+), annotated to this region with positions 100285632 and 100285650, respectively. Although these indel mutations may indeed occur at different positions, they are biologically equivalent because they result in the same altered sequence AGGAAAGAAAGAAAGAAAGAAAGAG. Since many databases such as dbSNP, Database of Genomic Variants (DGV), and Ensembl combine indels resulting from large-scale studies, similar cases often exist in those databases, leading to a nonnegligible problem of data redundancy. In fact, about 10% [4] of the human indels stored in dbSNP and 18% [1] in Ensembl are redundant. Resolving the indel redundancy in major databases is important for subsequent genetics research. Nevertheless, this problem has not been given the attention it deserves.

Numerous approaches have been developed for systematic comparison of indels to determine equivalence and hence solve the redundancy problem. The “strict matching” approach matches two indels if they share the same position, reference, and alternate alleles in two different entries in the VCF file. However, as demonstrated in [3], this approach fails to find equivalent indels that are not identical. The “distance based approach” treats two indels as equivalent if both have the same length and occur within a certain distance such as ± 5 bp [5] or ± 25 bp [6]. However, this approach introduces false positives when neighboring indels are not equivalent [1] and misses equivalent indels that are farther apart than the distance cutoff. Clearly, selection of an optimal distance cutoff is a tradeoff of the two types of errors: smaller distance cutoffs result in a decreased false positive rate but an increased false negative rate.

To address the limitations of the two aforementioned approaches, the more widely used “normalization” approach attempts to solve the indel redundancy problem by left (or right) normalization, i.e., consistently shifting the start position of an indel to the left (or right) as long as the resulting sequence is the same as the one generated by the original mutation [7]. Tools using this type of variant normalization include vt normalize [2], BCFtools [8], and GATK LeftAlignAndTrimVariants [9]. These tools usually take a VCF file as input, output another VCF file with canonical VCF entries for the indels after normalization, and then perform “strict matching” to find equivalent indels with exactly the same canonical representation. The normalization approach generally performs well in identifying equivalent indels, but as shown here, fails to normalize complex variants.

The positions of indels may get changed after left/right normalization, potentially misleading downstream analysis. For example, the deletion rs536379477 resides in the exon of the transcript ENST00000590192.1, but the equivalent deletion rs41436444 is in the intron of the same transcript. Therefore reporting these two indels with the same normalized position might lead to missing significant insight into genetic diseases or phenotypes of interest. Since the exact positions of most indel variations are not known, it is thus best to represent the indel of interest with a range of positions, within which equivalent indels can occur, rather than as a single normalized position. A similar idea was proposed by Krawitz et al. [10].

This paper proposes UPS-indel, a universal positioning system for indels, whereby every indel variant is represented by a range of positions within which all equivalent indels can occur. This representation is added to the VCF file resulting in a UVCF file containing not only the original indel calling results, but also the complete representation of all equivalent indels. The advantage of adding this column of information to the existing VCF file is (1) the original VCF file structure is unchanged so the UVCF file is still compatible with many downstream programs, (2) the UPS-indel notation facilitates the comparison of indels from different VCF files, (3) for equivalent indels that overlap both coding and noncoding regions, having the range column in the indel calling output would allow a downstream indel annotation system to consider the range rather than a single position, possibly annotating both a coding and noncoding variant. In summary, this work extends the previous work of Krawitz et al. [10] and Assmus et al. [1] by a new coordinate system Universal Positioning System (UPS), a rigorous mathematical proof that all (deletion and insertion) equivalent indels are found, the handling of complex variants, and a simple modification of an input VCF file to produce an output UVCF file containing the indel equivalence information. Results show that UPS-indel identifies more redundant indels than the existing approaches, also enables a comparison between indel calling results produced by different indel callers, and performs better than other state-of-the-art approaches for finding indels in common among call sets.

## Materials and Methods

This section defines some terms frequently used in this paper.

### Alternate Sequence

A sequence that is produced by introducing a specific indel to the reference sequence at a specific position. This is also known as the *mutant sequence*.

Let *R* be the reference sequence and *p* be either an insertion or a deletion of a given length that occurs at a given position in the reference sequence. The alternate sequence for insertion is denoted by *R′_1_*= *R* + *p* and for deletion by *R′_D_* = *R* – *p*.

### Equivalent Indels

Two indels are considered equivalent if and only if they produce the same alternate sequence. Note that equivalent indels must be of the same type (insertion and deletion) and same length.

### Redundant Indels

Equivalent indels that are reported as distinct entries in a VCF file are defined as redundant indels.

### Region of Equivalence

This is defined as the range of positions in the reference sequence where equivalent indels occur.

### Cyclic Permutation

A permutation (*y*_0_, *y*_1_, *y*_2_, …, *y_n_*_−1_) *f*(*x*_0_, *x*_1_, *x*_2_, …, *x_n_*_−1_) where y_i_ = *x*_(*i+k*)*mod n*_ for 0 ≤ *i* ≤ *n* − 1, *k* can be positive (left cyclic) or negative (right cyclic).

Table 1 shows an example of equivalent indels. Observe that all equivalent indels are cyclic permutations of each other (e.g., a cyclic permutation of CT is TC and cyclic permutations of TGT are GTT and TTG) and equivalence continues until there is a mismatch (see Supplementary Table 2). This observation leads to the following theorem.

**Table 1.** An example of equivalent indels.

| Equivalent insertions |  | Equivalent deletions |  |
| --- | --- | --- | --- |
| Reference, $R$ | GTCTA | Reference, $R$ | ACTGTTGTG |
| Case 1, $R'_I$ | G[TC/+]TCTA | Case 1, $R'_D$ | AC[TGT/-]TGTG |
| Case 2, $R'_I$ | GT[CT/+]CTA | Case 2, $R'_D$ | ACT[GTT/-]GTG |
| Case 3, $R'_I$ | GTC[TC/+]TA | Case 3, $R'_D$ | ACTG[TTG/-]TG |
| Case 4, $R'_I$ | GTCT[CT/+]A | Case 4, $R'_D$ | ACTGT[TGT/-]G |

#### Theorem 1

All equivalent indels in the region of equivalence are cyclic permutations of each other.

**Proof:** Consider two equivalent indels *d_1_* and *d_2_* and the equivalence region *R* they define. For insertion within *R*, the alternate sequences are

*d_1_S = Sd_2_*

for some nonempty *S.* For deletion within *R*, the alternate (possibly empty) sequence is *S* starting with

*d_1_S = Sd_2_*.

**Case 1.** For |*S*| < |*d_1_*|, *d_1_* = *SX* for nonempty *X* and *d_1_S* = *SXS* = *Sd_2_* implies *d_2_* = *XS* is a cyclic permutation of *d_1_* = *SX*.

**Case 2**. For |*S*| = |*d_1_*|, *d_1_ = d_2_ = S*.

**Case 3.** For |*S*| > |*d_1_*|, *S* = *d_1_X* for nonempty *X* with |*X*| < |*S*|, and *d_1_d_1_X* = *d_1_S = Sd_2_* = *d_1_Xd2* implies *d_1_X* = *Xd_2_*. Repeating this argument for *d_1_X* = *Xd_2_* eventually reduces *X* to one of the previous two cases.

Another case for deletion is when *R* is periodic with period |*d_1_*|, having the form *R* = *d_1_ d_1_*…….. *d_1_* (*d_1_*)_1_ where (*d_1_*)_1_ is the first symbol of *d_1_*. Then every consecutive subsequence *d_2_* of *R* with |*d_1_*| = |*d_2_*| is an equivalent deletion, and *d_2_* is a cyclic permutation of *d_1_*. **(Q.E.D)**

**Corollary.** For |*S*| > |*d_1_*|, *S* must have the form *d_1_ d_1_*…….*…… *d_2_ d_2_* with an equal number of *d_1_*s and *d_2_*s.

Based on the theorem, an algorithm called UPS-indel (see Table 2) exhaustively increases the range of equivalence as far as possible in both left and right directions from a given indel position. Finally for each indel in the VCF file, the algorithm reports its range of equivalence, which is called the Universal Positioning System coordinate (UPS-coordinate). Once indels are represented by their UPS-coordinates, identifying redundant indels becomes a trivial task of string comparison (e.g., Fig. 2(A), comparison across the 8^th^ column). Note that since UPS-indel implements Theorem 1, which characterizes indels within an equivalence region, UPS-indel is exhaustive, finding all equivalent indels.

**Figure 1:**
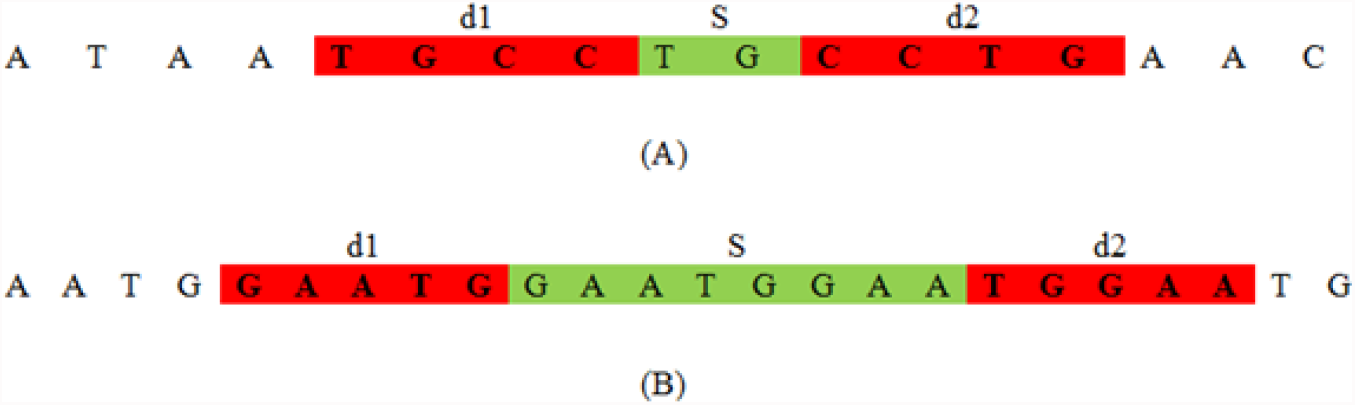
Illustration of two cases of Theorem 1. (A): |d_1_| > |S|, (B): |d_1_| < |S|.

**Figure 2.**
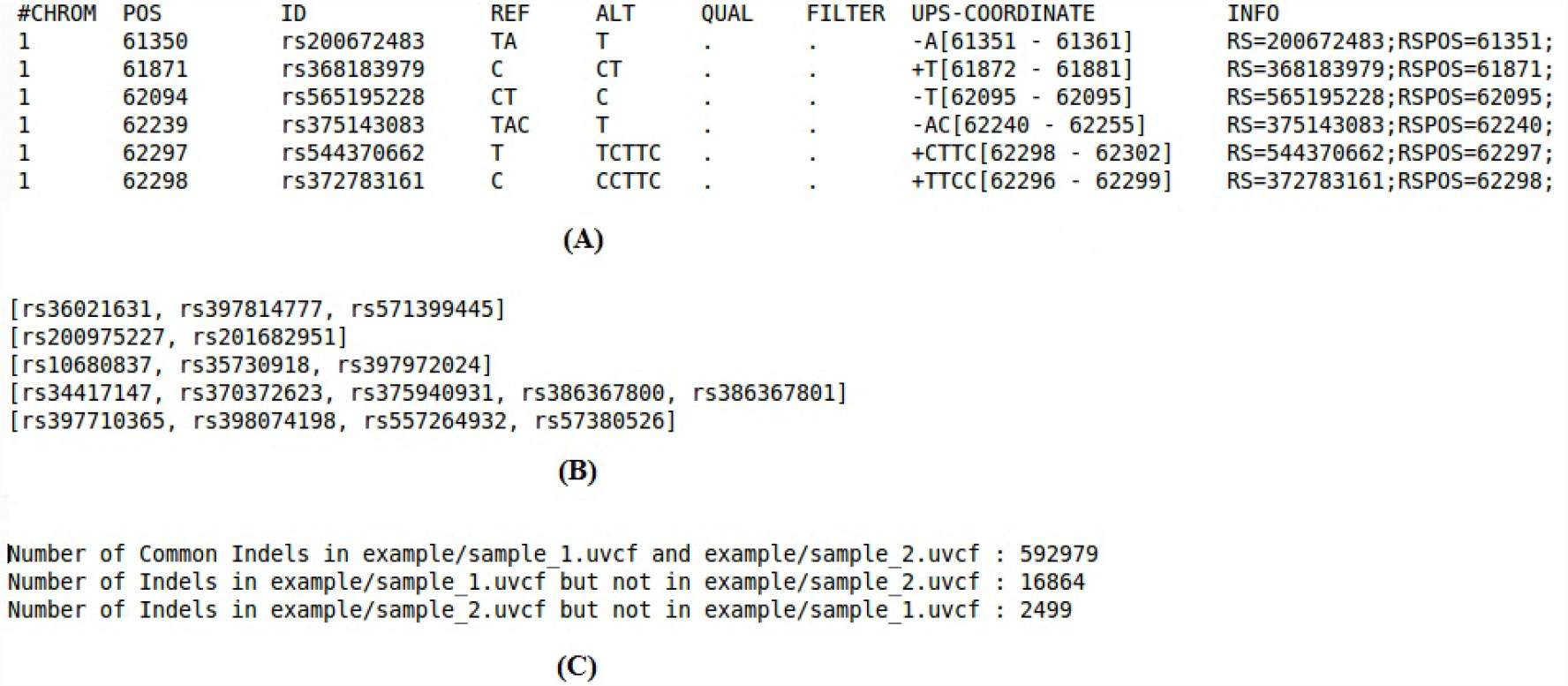
Different utilities of UPS-indel. (A)UVCF format, (B) redundant indel list, and (C) comparing two uvcf files.

**Table 2.**
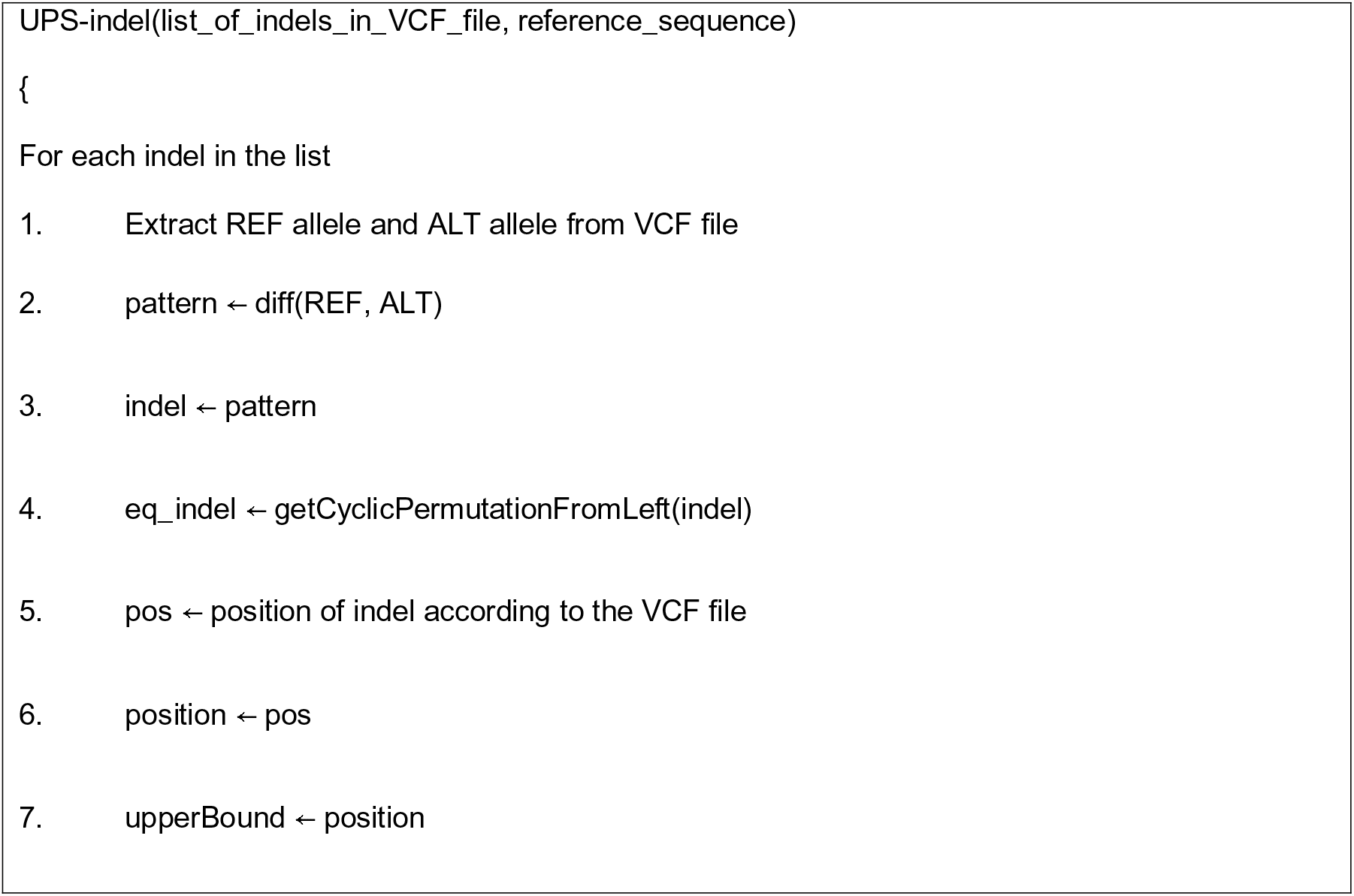
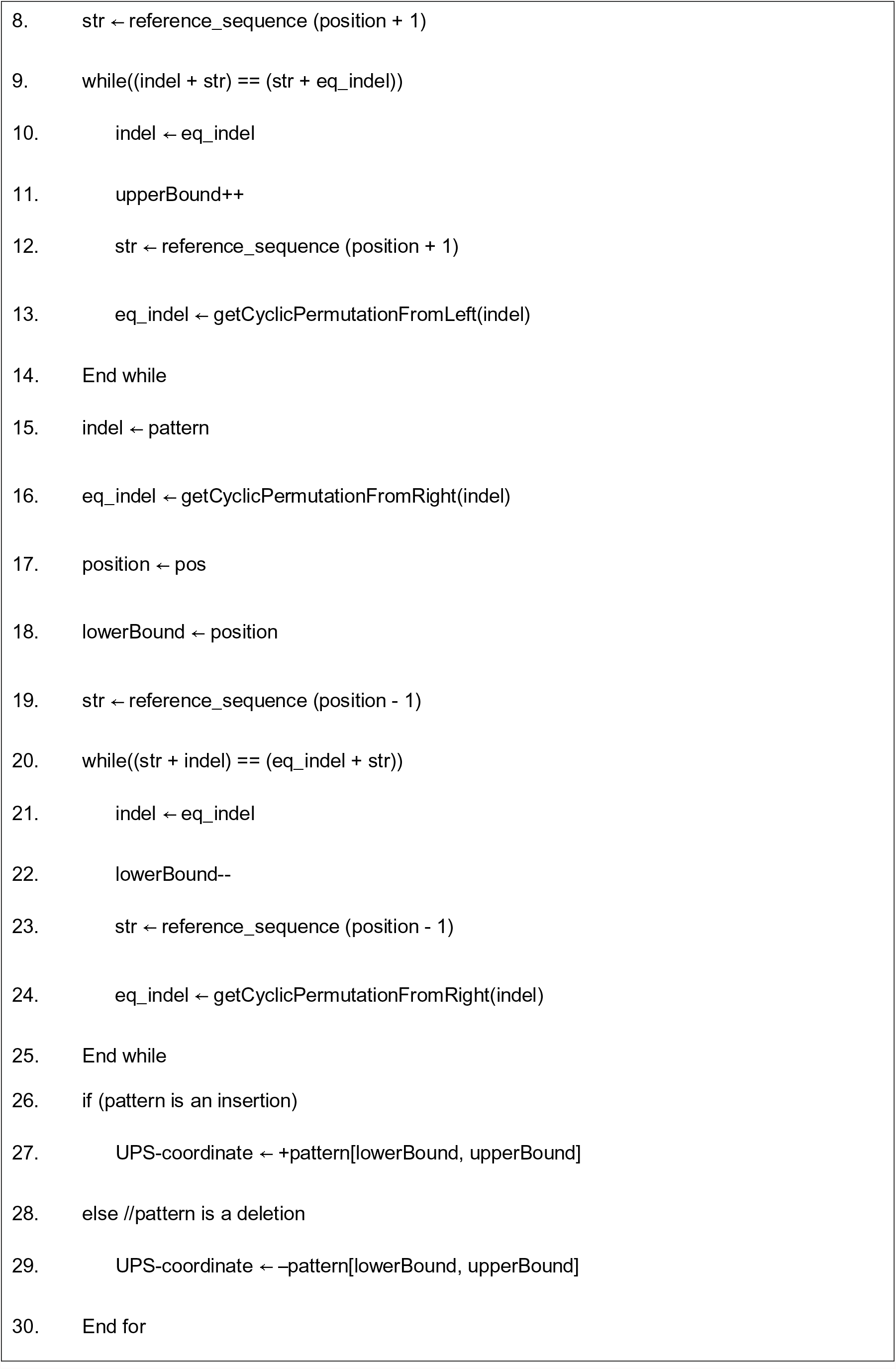
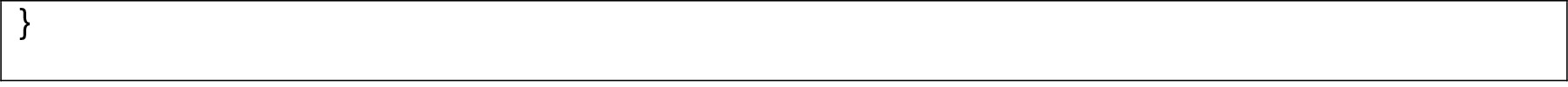
UPS-indel algorithm.

Note that “left” and “right” cyclic permutations are equivalent – there is no difference. In line 2 of the UPS-indel algorithm, while extracting the “pattern” from the entries of the RFE and ALT columns of the input VCF file, UPS-indel performs horizontal decompositions of the complex variants and assigns the indel part as the value of pattern. For example, suppose in the REF column of a VCF entry there is an allele “ATAA” and in the ALT column there is an allele “AG”. In this case, UPS-indel performs horizontal decompositions of the complex variants to produce two separate entries (AT → AG and AA →<empty> meaning that there is a deletion of AA).

UPS-indel is written in C++ and can run on Linux, Windows, or Mac operating systems that have a C++ compiler. UPS-indel uses SeqAn, an open source C++ library containing efficient algorithms and data structures to analyze large genome sequences [11]. The input of UPS-indel is a reference chromosome sequence, a VCF file containing a list of indels, and an output file name, for example,

~~~
./ups_indel example/chr1.fa example/chr1.vcf example/chr1.uvcf
~~~

This command line produces an output file named chr1.uvcf, containing the UPS-coordinates of all the indels in chr1.vcf. Figure 2(A) shows an example UVCF file.

The UVCF file keeps the same content/format as the VCF file, with an additional column that contains the indel's UPS-coordinate information. The interpretation of the UPS-coordinate follows:

- Symbols + and – denote insertion and deletion, respectively, followed by the base pairs inserted/deleted from the reference, and the UPS-coordinate (in square brackets).
- The UPS-coordinate contains a range of positions in the square brackets representing the region of equivalence for the indel. For example, the UPS-coordinate + CTTC [62298 - 62302] means there is an insertion of CTTC at position 62298, and the same alternate sequence can be produced by inserting TTCC at position 62299, or TCCT at position 62300, and so on.

Once indels are represented by the coordinates produced by UPS-indel, one can easily identify redundant indels within one indel call set or multiple indel call sets. For example, the following command line

~~~
./ups_generate_redundant_indel_list example/chr1.uvcf example/redundant_indel_list.txt
~~~

produces a list of indel groups containing dbSNP IDs of redundant indels (Figure 2(B)).

UPS-indel groups all redundant indels together. For example, consider a group [rs34748242, rs59148039] with the UVCF entry shown in Table 3. These two indels belong to the same indel type (insertion), have same base pairs inserted (TG), and share the same UPS-coordinate and hence they are considered as equivalent.

**Table 3.** UVCF file for redundant indels.

| #CHRM | POS | ID | REF | ALT | QUAL | FILTER | UPS-COORDINATE |
| --- | --- | --- | --- | --- | --- | --- | --- |
| 1 | 10009638 | rs34748242 | T | TTG | . | . | +TG[10009639 - 10009648] |
| 1 | 10009639 | rs59148039 | T | TGT | . | . | +TG[10009639 - 10009648] |

UPS-indel can compare multiple indel call sets. This utility is particularly useful for generating a high-confidence indel call set by taking the intersection of the results of different indel callers [12], or merging the indel calling results from different tools for a consensus variant caller [13], or comparing indel call sets generated by different indel callers to determine their relative recall, precision, and accuracy, and to understand the source of their dissimilarities. To use this utility of UPS-indel, after converting two VCF files to UVCF files, one can use the following command to get the comparison result (Figure 2(C)), which contains useful statistics for downstream analysis:

~~~
./ups_compare_uvcf_files example/sample1.uvcf example/sample2.uvcf example/comparison_result.txt
~~~

All of the above mentioned utilities of UPS-indel are also available at http://bench.cs.vt.edu/ups-indel/ (Figure 3).

UPS-indel is compared with other existing tools that also find equivalent indels through variant normalization. These tools include vt normalize (version 0.5) [2], BCFtools (version 1.3) [8], and GATK LeftAlignAndTrimVariants (version 3.5) [9]. Like UPS-indel, all of these tools take a VCF file and the reference genome as input and produce the normalized position of the indels in the VCF file.

**Figure 3.**
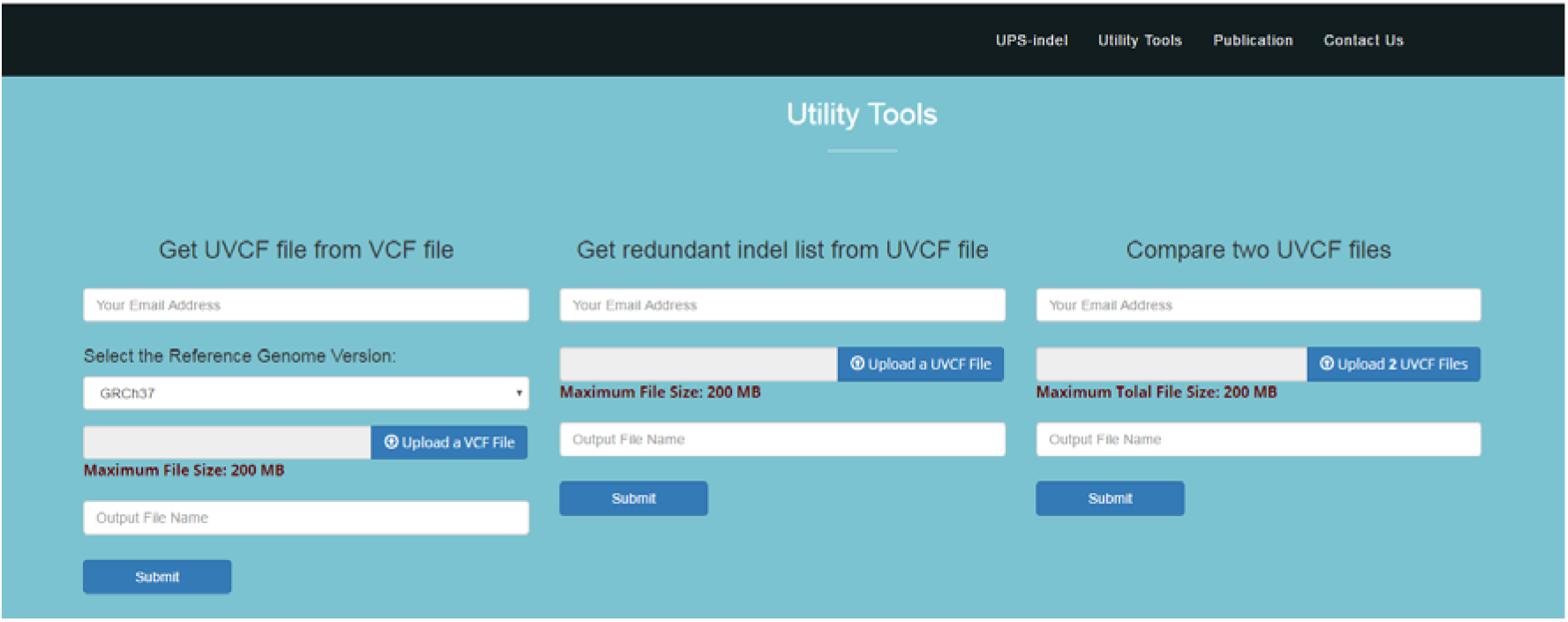
Main user interface of UPS-indel.

Another tool Vindel [4] also finds equivalent indels using a heuristic approach, but was not included in the comparison as it uses a flat file as input instead of a VCF file.

A VCF file of dbSNP (version 142, GRCh37p13) and the GRCh37 reference genome were used as the inputs to these tools. The VCF file contains both SNPs and indels, and VCFTools [14] is used to extract indels from the VCF file. The comparison was extended to the COSMIC dataset as well.

There are other tools that could also be considered for comparison. Both VarMatch [3] and RTGTools [15] use a branch and bound algorithm to search for equivalent indels. They are not suitable for processing population scale indel call sets such as dbSNP and COSMIC because densely packed indels in such datasets make the search space too large to be processed by a branch and bound algorithm. READDI [16] considers repeat-induced ambiguities as well as tool-induced inaccuracies while searching for equivalent deletions using the longest common extension algorithm. This tool is limited to finding deletions only, and hence not included in the comparison for the dbSNP and COSMIC datasets. Nevertheless, in this study a smaller dataset is used to compare UPS-indel with VarMatch (Version available on April 5, 2017), RTGTools (Version 3.7.1), and READDI (Version available on April 5, 2017).

## Results and Discussion

### Finding equivalent indels in the dbSNP dataset

The input VCF file contains about 8.9 million indels from the human genome. For this input, UPS-indel produces the UVCF file and the other three tools, vt normalize, BCFtools, and GATK LeftAlignAndTrimVariants, generate the normalized VCF file. These three tools perform left normalization of indels and output a left normalized representation. Therefore, for these three tools, two indels are equivalent if and only if they satisfy the following conditions:

1. Both indels are of the same type (insertion or deletion).
2. Both indels share the same pattern after normalization: [value of the REF column in the normalized VCF file – value of the ALT column in the normalized VCF file – value of the POS column in the normalized VCF file]. Note that one might think that considering the position should suffice, because after normalization, equivalent indels should have the same position in the VCF file. However, the example in Table 4 shows that indels rs371246544 and rs71724031 have the same normalized position but are not equivalent.

**Table 4.**
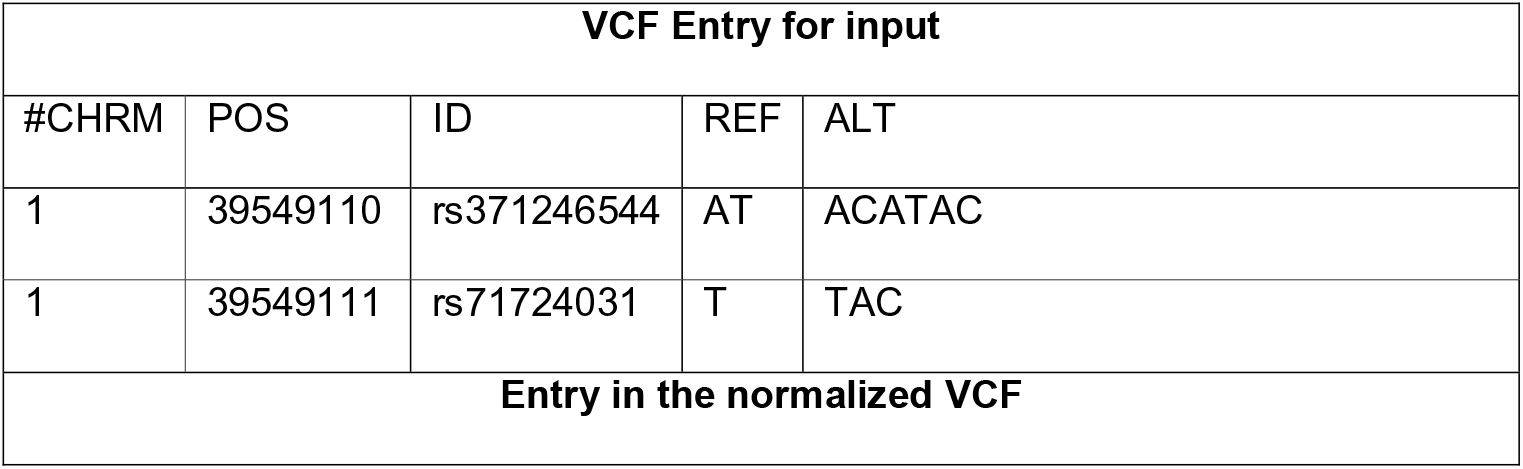
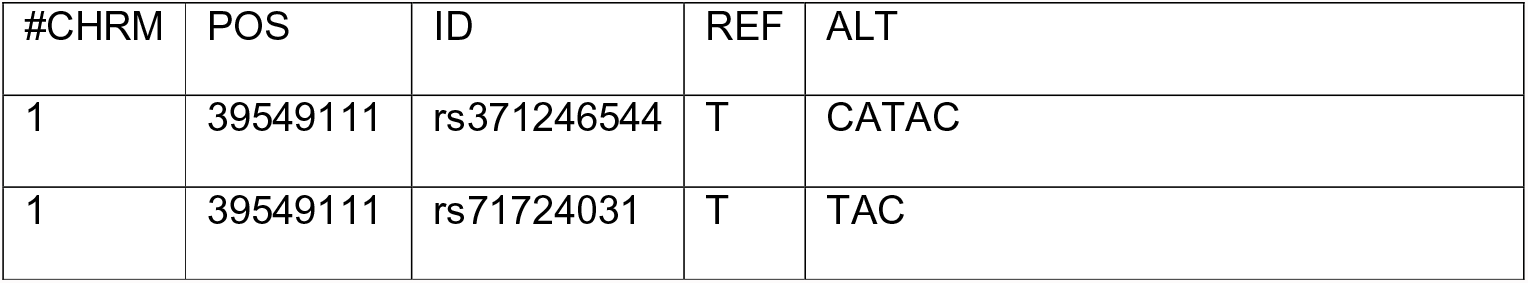
An example explaining why considering only normalized position does not suffice for identifying redundant indels for vt normalize and BCFtools.

| VCF Entry for input |  |  |  |  |
| --- | --- | --- | --- | --- |
| #CHRM | POS | ID | REF | ALT |
| 1 | 39549110 | rs371246544 | AT | ACATAC |
| 1 | 39549111 | rs71724031 | T | TAC |
| Entry in the normalized VCF |  |  |  |  |

| #CHRM | POS | ID | REF | ALT |
| --- | --- | --- | --- | --- |
| 1 | 39549111 | rs371246544 | T | CATAC |
| 1 | 39549111 | rs71724031 | T | TAC |

The comparison is based on the criterion: the redundant indel ratio =

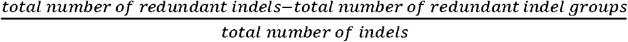

where the numerator is the total number of redundant indels reported since only one indel from each redundant indel group should be reported in the output and the remaining should be considered as redundant.

Figure 4 shows the comparison of the redundant indel ratios reported by UPS-indel, vt normalize, BCFtools, and GATK LeftAlignAndTrimVariants for indels in the dbSNP dataset. For the entire human genome, UPS-indel identified ∼ 15% redundant indels (see Supplementary Table 3 and Supplementary Figure 1 for chromosome-wise comparison), as compared to 11.82% by vt normalize, 11.82% by BCFtools, and 11.81% by GATK LeftAlignAndTrimVariants. At the chromosome level, UPS-indel identified about 3% more redundant indels than the other three tools.

**Figure 4.**
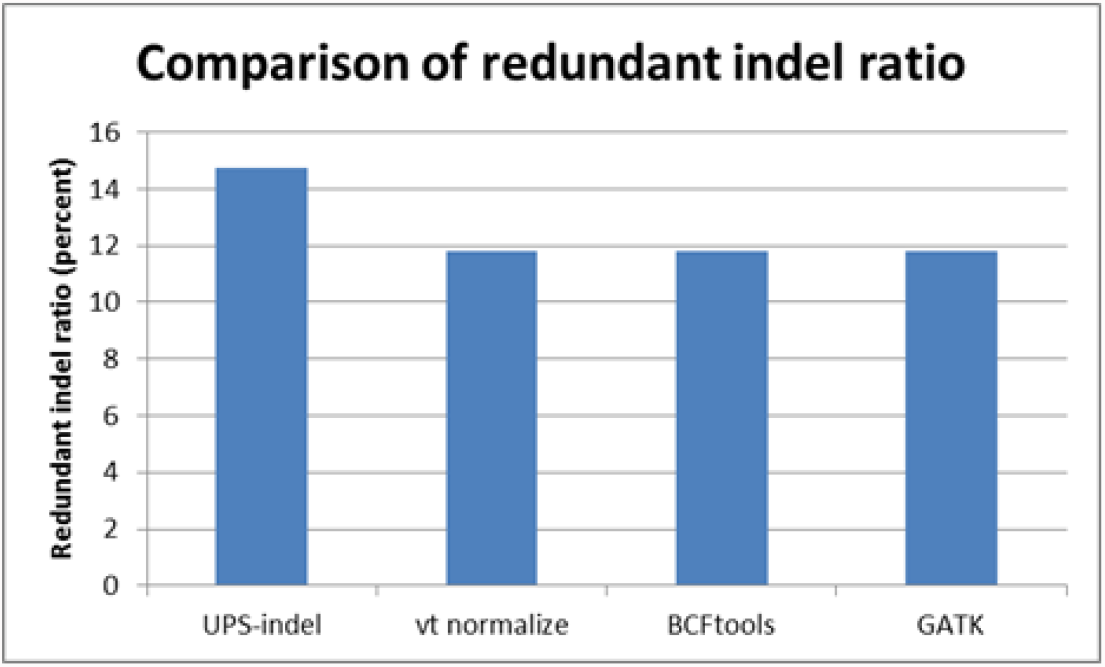
Comparison among the tools based on redundant indel ratio for the dbSNP dataset.

Examining the sets of redundant indels detected by UPS-indel and the other tools shows that vt normalize and BCFtools produce exactly the same results for all chromosomes. Moreover, all the redundant indels detected by vt normalize, BCFtools, and GATK LeftAlignAndTrimVariants are also detected by UPS-indel, as shown in Figure 5. Further, for all chromosomes, UPS-indel identified a total of 456,352 more redundant indels than the other tools. As proved in the methods, UPS-indel identifies all the redundant indels, the comparison result shows that the other three tools are not exhaustive in finding all the redundant indels.

**Figure 5.**
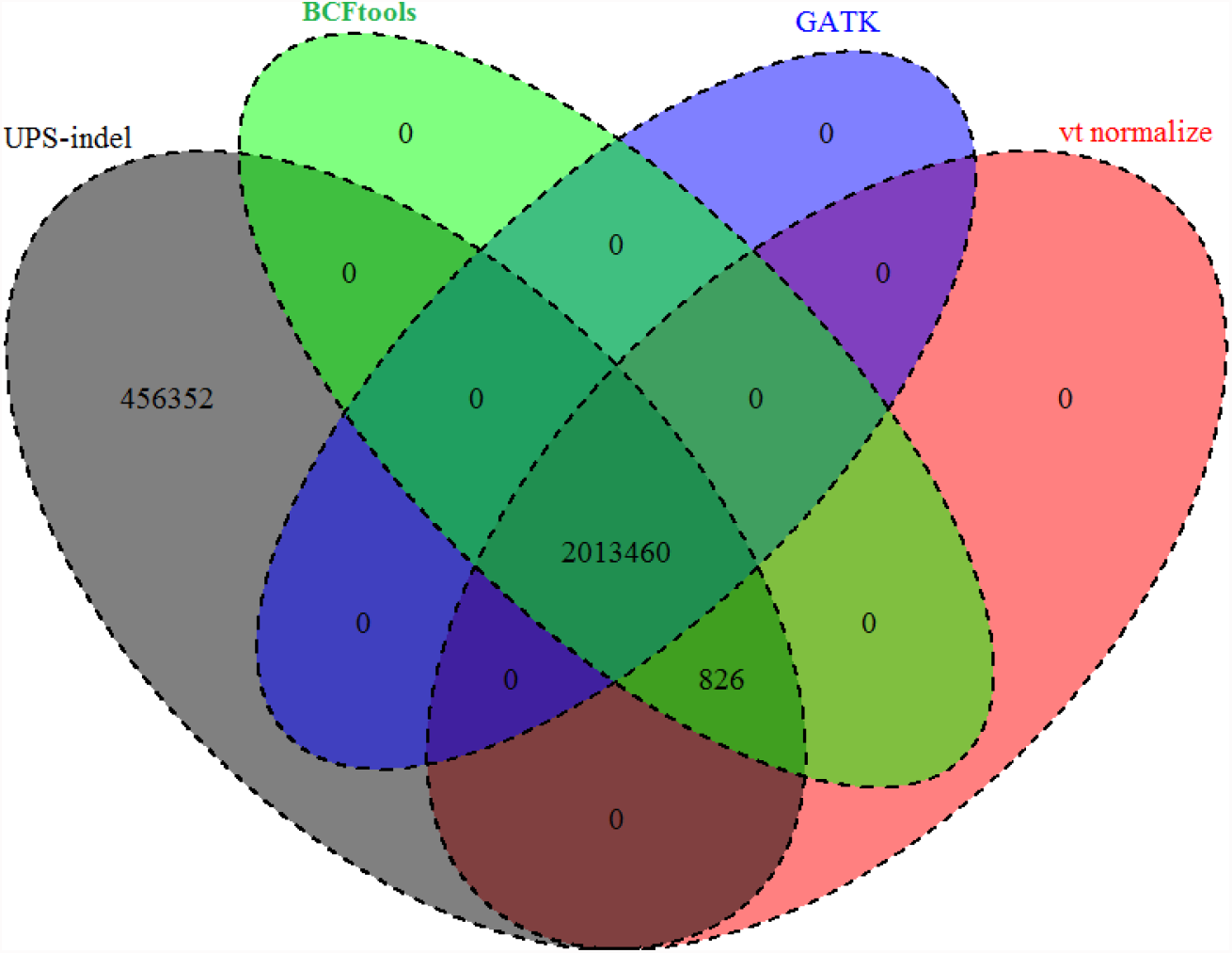
Venn diagram to compare the number of redundant indels detected by UPS-indel and other tools. (Venn Diagrams are generated using the R package VennDiagram [1 7].)

Why are several indels found as redundant by UPS-indel but not by other tools? An investigation shows that these equivalent indels are missed by the other tools because, due to the computation time limit, they cannot exhaustively search every cyclic permutation at every feasible position as is done by UPS-indel. For example, long multiallelic indels are not considered by default for normalization. Had the tools considered these indels separately, they would have been able to find an equivalent indel located at a different position. For this situation, UPS-indel splits the VCF entry into multiple entries by default and considers each of the indels separately while finding redundant indels. Table 5 provides such an example.

**Table 5:**
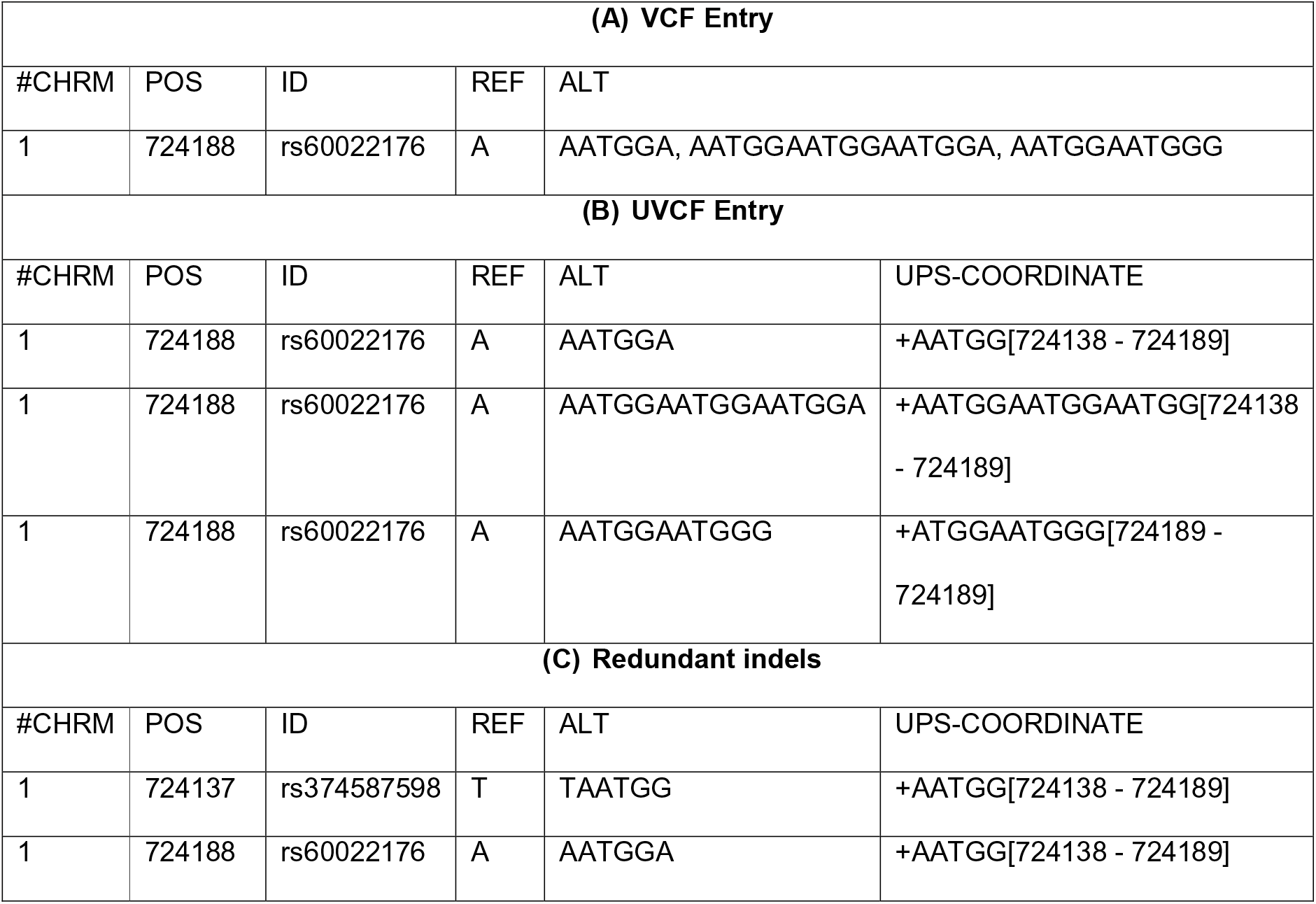
Example of multiallelic insertion type indels missed by other tools but detected as redundant by UPS-indel.

For the indel shown in Table 5 (panel A), no normalization was done by vt normalize, BCFtools, or GATK LeftAlignAndTrimVariants. UPS-indel splits the entry into three indels and finds the UPS-coordinate for each of them separately (Table 5, panel B). Splitting the VCF entry and considering the indels separately, UPS-indel managed to find another indel equivalent to one of the indels (Table 5, panel C). Therefore UPS-indel reports indels with id rs374587598 and rs60022176 as redundant.

The example in Table 5 is for insertion; an example for deletion is illustrated in Table 6.

**Table 6.** Example of multiallelic deletion type indels missed by other tools but detected as redundant by UPS-indel.

| VCF Entry |  |  |  |  |  |
| --- | --- | --- | --- | --- | --- |
| #CHRM | POS | ID | REF | ALT |  |
| 1 | 7552657 | rs376707888 | GTG | G, GTGCA |  |
| UVCF Entry |  |  |  |  |  |
| #CHRM | POS | ID | REF | ALT | UPS-COORDINATE |
| 1 | 7552657 | rs376707888 | GTG | G | -GT[7552657 - 7552658] |
| 1 | 7552657 | rs376707888 | GTG | GTGCA | +CA[7552658 - 7552658] |
| **Redundant indels** |  |  |  |  |  |
| #CHRM | POS | ID | REF | ALT | UPS-COORDINATE |
| 1 | 7552656 | rs139294420 | CGT | C | -GT[7552657 - 7552658] |
| 1 | 7552657 | rs376707888 | GTG | G | -GT[7552657 - 7552658] |

In addition to the scenario mentioned above, GATK LeftAlignAndTrimVariants does not normalize any of the multiallelic indels regardless of the size which is also mentioned in [2]. Table 7 shows an example of this occurrence explaining why GATKLeftAlignAndTrim finds fewer number of redundant indels than vt normalize and BCFtools.

**Table 7.** Example of a multiallelic indel that is normalized by vt normalize and BCFtools but not by GATKLeftAlignAndTrim.

| **VCF Entry for dbSNP** |  |  |  |  |
| --- | --- | --- | --- | --- |
| #CHRM | POS | ID | REF | ALT |
| 1 | 823905 | rs397728418 | AA | A, AAA |
| **VCF Entry for GATK LeftAlignAndTrimVariants** |  |  |  |  |
| #CHRM | POS | ID | REF | ALT |
| 1 | 823905 | rs397728418 | AA | A, AAA |
| **VCF Entry for vt normalize and BCFtools** |  |  |  |  |
| #CHRM | POS | ID | REF | ALT |
| 1 | 823903 | rs397728418 | GA | G, GAA |

One might think that decomposing multiallelic indels into several biallelic indels produces the same results as UPS-indel for the normalization tools. To check this, the “decompose” utility of vt was used to perform a vertical decomposition of multiallelic indels into biallelic indels. Applying vt normalize to the decomposed indels could not find equivalent indels for complex variants, whereas UPS-indel is able to find the equivalent indels. Table 8 shows an example of this occurrence. Since vt normalize and BCFtools produce exactly the same results, these complex variants are missed by BCFtools as well.

**Table 8.** Example of a complex variant that is missed by vt normalize but detected as redundant by UPS-indel.

| VCF Entry (A) |  |  |  |  |  |
| --- | --- | --- | --- | --- | --- |
| #CHRM | POS | ID | REF | ALT |  |
| 1 | 2273131 | rs369694942 | GAAA | G |  |
| 1 | 2273140 | rs373243812 | AAAAA | AG |  |
| UVCF Entry (B) |  |  |  |  |  |
| #CHRM | POS | ID | REF | ALT | UPS-COORDINATE |
| 1 | 2273131 | rs369694942 | GAAA | G | -AAA[2273132 - 2273147] |
| 1 | 2273140 | rs373243812 | AAAAA | AG | -AAA[2273132 - 2273147] |

In the example shown in Table 8, VCF entries for the indels with ids rs369694942 and rs373243812 remain the same in the input and the output for vt normalize (Panel A), i.e., no normalization is done. Here the second indel (rs373243812) is a complex variant containing both aSNP (A → G) and a deletion of length three (AAA), and is ignored by vt normalize. However, UPS-indel performs a horizontal decomposition of the complex variant to produce two separate entries (AA→AG and AAA→<empty>) and finds the equivalent indel with id rs369694942 having a deletion of length three (AAA) in the UPS-Coordinate 2273132 to 2273147 (Panel B).

## Finding equivalent indels in the COSMIC dataset

UPS-indel was used to find redundant indels in the COSMIC (Catalogue Of Somatic Mutations In Cancer) dataset, the world's most comprehensive resource for exploring the impact of somatic mutations in human cancer [18]. With data collected for more than 2,500 human cancers, this archive describes millions of coding mutations, noncoding mutations, and other gene expression variants across the human genome.

For all chromosomes in the COSMIC dataset, UPS-indel identified 28.17% and 13.11% redundant indels in the COSMIC coding and noncoding indel datasets, respectively, which are higher than the redundant indel ratios reported by the other tools. Figure 6 shows the comparison of the redundant indel ratios reported by UPS-indel, vt normalize, BCFtools, and GATK LeftAlignAndTrimVariants for both the COSMIC coding and noncoding datasets. Comparisons for chromosome-wise redundant indel ratios among the tools are given in Supplementary Materials (See Table 4 and Figure 2 for COSMIC coding and Table 5 and Figure 3 for noncoding indels).

**Figure 6.**
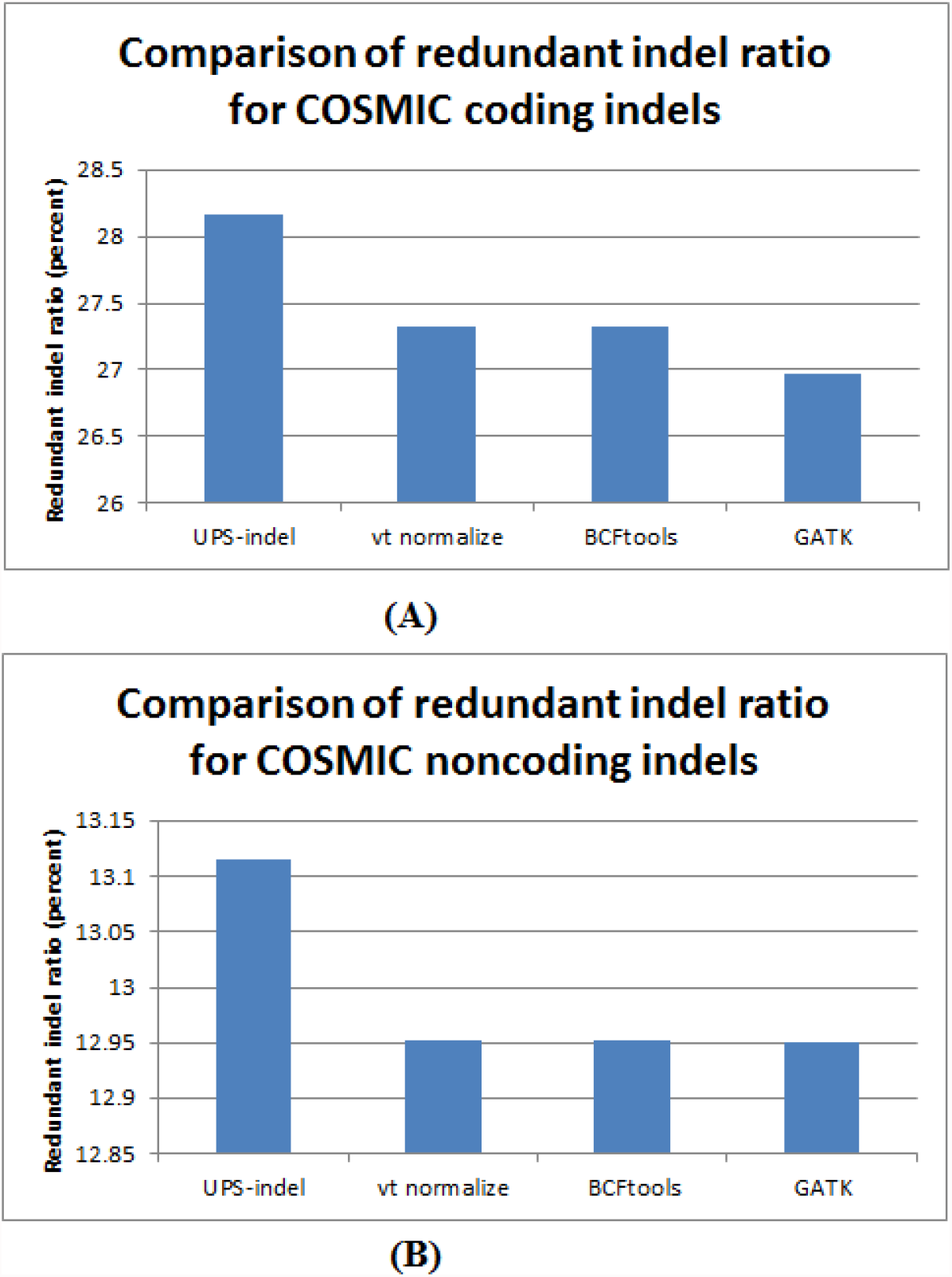
Comparison of redundant indel ratio for (A) COSMIC coding and (B) COSMIC noncoding indels.

Similarly, examining the sets of redundant indels identified by the tools, Figure 7 shows that for both the COSMIC coding and noncoding indels, UPS-indel identified all the redundant indels detected by the other tools. In addition to that, for the whole genome, 2,118 (Figure 6A) and 553 (Figure 6B) unique redundant indels for COSMIC coding and noncoding indels, respectively, are detected by UPS-indel but missed by other tools.

**Figure 7.**
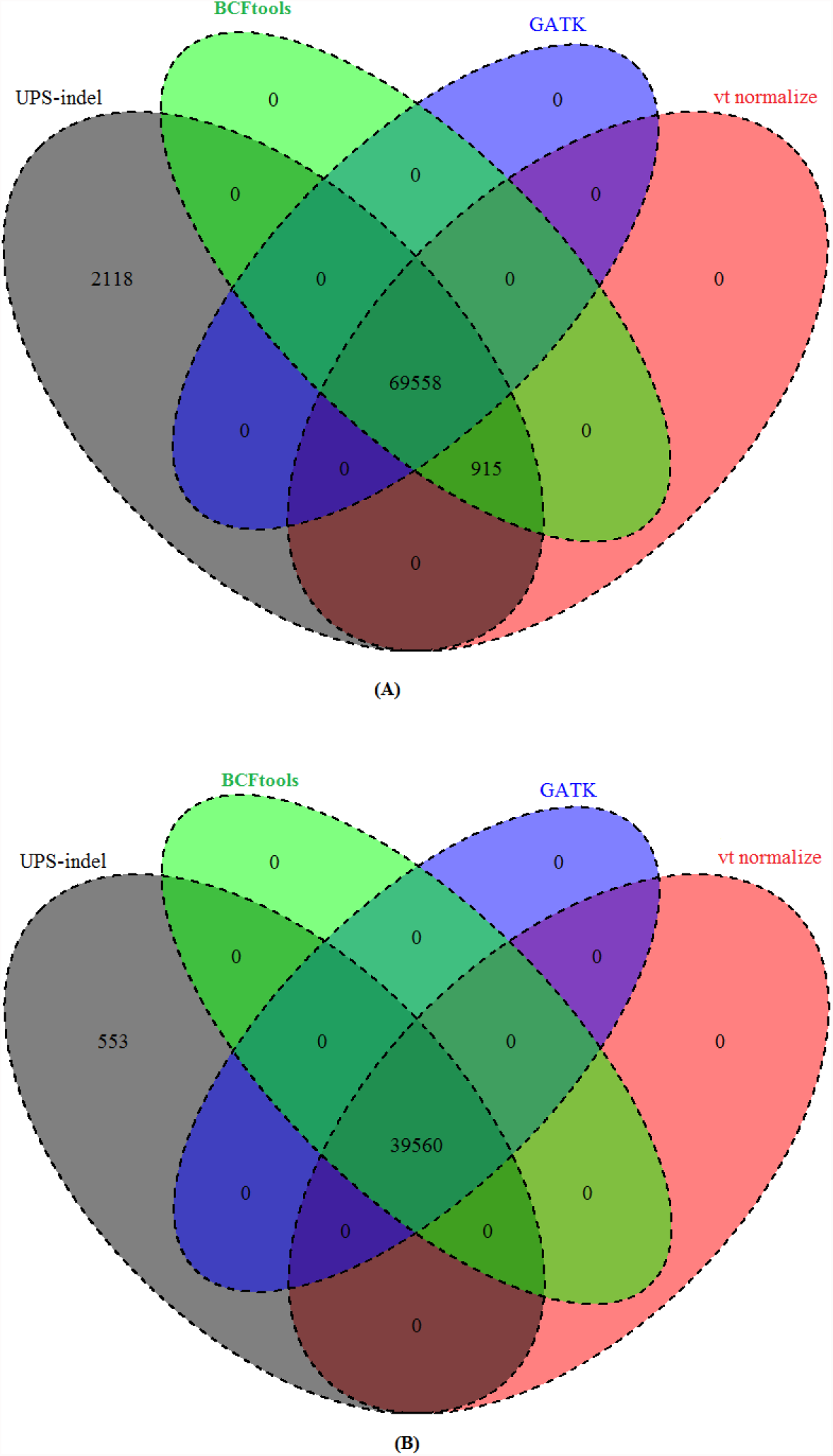
Venn diagram to compare the number of redundant indels detected by UPS-indel and other tools in (A) COSMIC coding and (B) COSMIC noncoding indel datasets.

As for dbSNP, the reason why some COSMIC coding and noncoding indels were considered as redundant by UPS-indel but missed by other tools is that, in the normalized VCF for these other tools, redundant indels must contain the same pattern: [value of the REF column in the normalized VCF file – value of the ALT column in the normalized VCF file – value of the POS column in the normalized VCF file]. The reason for this pattern match restriction was given earlier. In Table 9, all tools except UPS-indel missed the indel with id COSM5068028 in the redundant indel group consisting of indels with id COSM3732389 and id COSM5348791, because of not having the same pattern. Therefore it might be assumed that only normalized position should be considered to group them together. However, then the indel with id COSM3685916 would be placed in the same group, although it is a deletion type indel whereas the others are insertion type indels, and also the resultant sequences are different. UPS-indel groups the indels correctly by placing indels with ids COSM5068028, COSM3732389, and COSM5348791 in the same redundant indel group as they have the same base pair inserted, have the same region of equivalence, and also are of the same indel type.

**Table 9.**
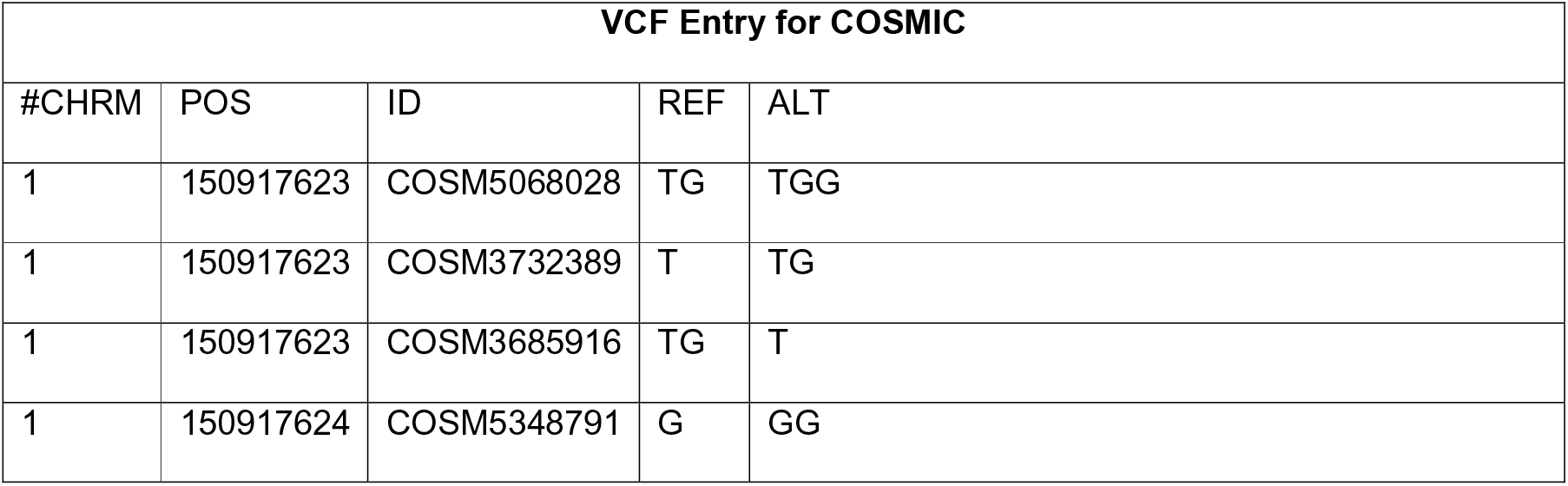
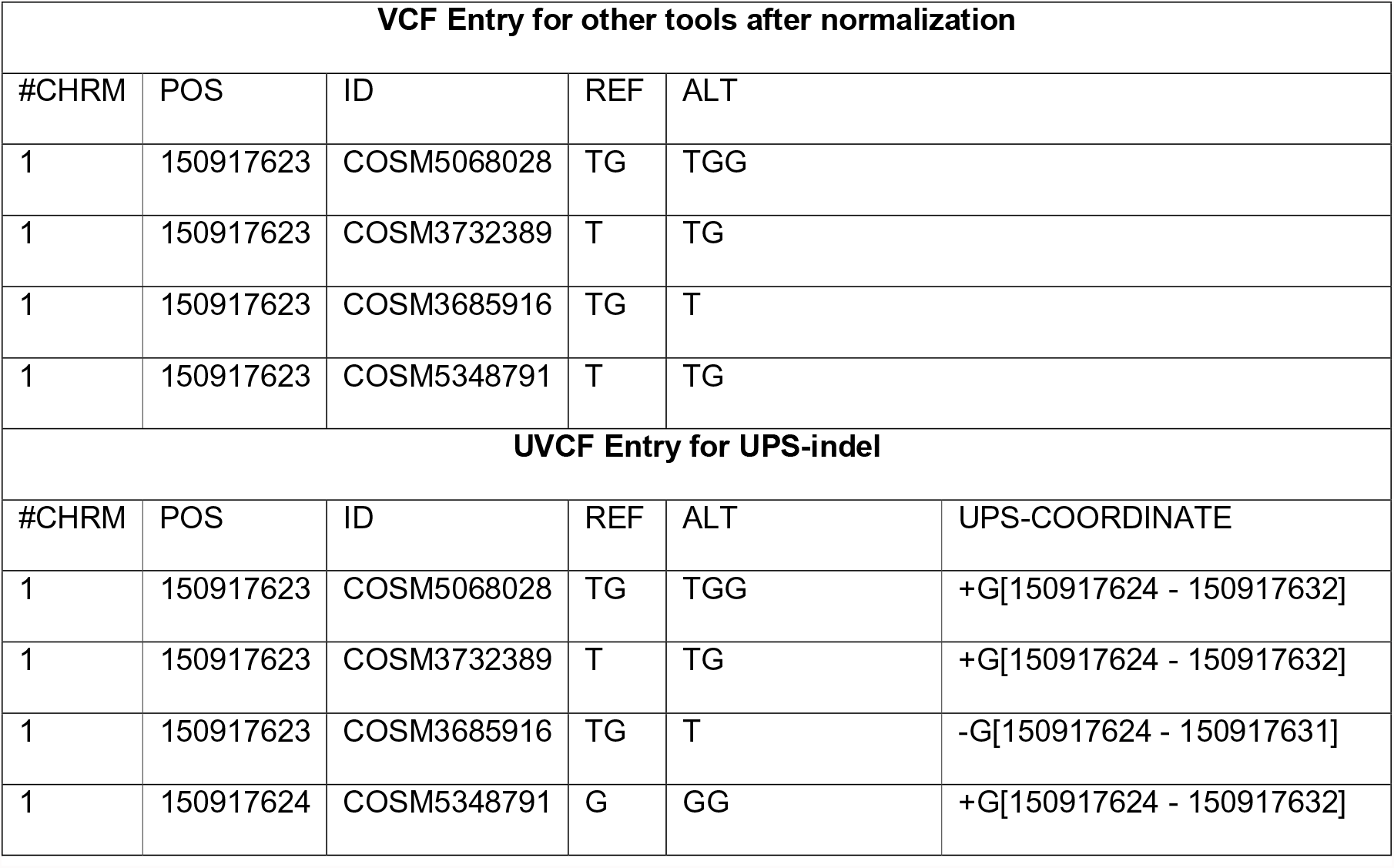
Example of COSMIC indel that is missed by other tools but detected as redundant by UPS-indel.

| VCF Entry for COSMIC |  |  |  |  |
| --- | --- | --- | --- | --- |
| #CHRM | POS | ID | REF | ALT |
| 1 | 150917623 | COSM5068028 | TG | TGG |
| 1 | 150917623 | COSM3732389 | T | TG |
| 1 | 150917623 | COSM3685916 | TG | T |
| 1 | 150917624 | COSM5348791 | G | GG |

| VCF Entry for other tools after normalization |  |  |  |  |  |
| --- | --- | --- | --- | --- | --- |
| #CHRM | POS | ID | REF | ALT |  |
| 1 | 150917623 | COSM5068028 | TG | TGG |  |
| 1 | 150917623 | COSM3732389 | T | TG |  |
| 1 | 150917623 | COSM3685916 | TG | T |  |
| 1 | 150917623 | COSM5348791 | T | TG |  |
| UVCF Entry for UPS-indel |  |  |  |  |  |
| #CHRM | POS | ID | REF | ALT | UPS-COORDINATE |
| 1 | 150917623 | COSM5068028 | TG | TGG | +G[150917624 - 150917632] |
| 1 | 150917623 | COSM3732389 | T | TG | +G[150917624 - 150917632] |
| 1 | 150917623 | COSM3685916 | TG | T | -G[150917624 - 150917631] |
| 1 | 150917624 | COSM5348791 | G | GG | +G[150917624 - 150917632] |

GATKLeftAlignAndTrimVariants found fewer redundant indels than other tools because GATKLeftAlignAndTrimVariants does not consider very large indels for normalization. For example, the indels with ids COSM5196837 and COSM5066846, which are deletions of length 371bps and 222 bps, respectively, are not considered by GATKLeftAlignAndTrimVariants for normalization. The reason is that GATK LeftAlignAndTrimVariants uses 200 bps as the default size of the sliding window on the reference (the parameter `--reference_window_stop`) while left aligning the alleles which is smaller than the length of the missed deletions.

These tools are also compared based on the average running time taken to process the input VCF file for normalization (by vt normalize, BCFtools, and GATKLeftAlignAndTrimVariants) or for generating the UPS-Coordinate (by UPS-indel). All tools were run on a desktop computer having an Intel Core i7-2600 CPU with eight cores (at 3.40 GHz) and 16GB of RAM. Table 10 shows the average running time for chromosome 1 of the dbSNP VCF file. Among these tools BCFtools is the fastest taking 6 seconds followed by vt normalize (6.18 seconds), GATK LeftAlignAndTrimVariants (17.22 seconds), and UPS-indel (35.22 seconds), which is the slowest. Since UPS-indel searches for equivalent indels exhaustively and is theoretically rigorous, the computation time is not surprisingly higher than that for other heuristic normalization tools.

**Table 10.** Comparison between VarMatch, RTG Tools, READDI, and UPS-indel of the number of overlaps found between the baseline and query call sets from chromosome 11 of an individual.

| Variant Caller Name | VarMatch (EVQ mode) | RTG Tools | READDI | UPS-indel |
| --- | --- | --- | --- | --- |
| Dindel | 8,933 | 8,717 | 8,796 | 8,973 |
| GATK Haplotype Caller | 11,113 | 10,765 | 10,954 | 11,129 |
| GATK Unified Genotyper | 7,734 | 7,596 | 7,563 | 7,734 |
| Pindel | 6,507 | 3,654 | 9,524 | 9,836 |

## Evaluating UPS-indel's performance in comparing different indel call sets

In genomic research related to indel calling, an important step in downstream analysis is to compare multiple indel call sets for (1) generating a highly accurate benchmark indel call set by taking the intersection of multiple call sets as done by Zook et al. [12] for the sample NA12878, (2) merging the call sets of different indel callers in a consensus caller as done by Trubetskoy et al. [13] for exome data, and (3) evaluating the accuracy of a newly proposed indel calling tool by comparing its indel call set with the benchmark call set. Comparing different indel call sets is also a common step in studies comparing the performance of different indel callers as done in [5],[19], and [20]. Different indel callers having different representations of the same indel complicates the comparison of different indel call sets. In addition to strict matching of indels, as mentioned earlier, a naïve but previously commonly used approach to compare multiple indel calling results is based on a simple distance criterion, that is, indels are considered to be equivalent if they are within a distance threshold (e.g., ±5 bp or ± 25 bp). For example, the original 1000 Genomes project used ± 25bp to compare multiple indel calling results [6]. To illustrate the advantage of using a UVCF file instead of a distance criterion or normalized VCF for comparing multiple VCF files, the alignment file for chromosome 11 of a single sample (HG00851) was picked up from the 1000 Genomes project and five indel callers: Dindel [21], GATK Unified Genotyper [9], GATK Haplotype Caller, Platypus [22], and Pindel [23] were used to produce VCF files for indels. The resultant VCF files were compared to determine the number of common indels from these five tools using three different approaches, namely a distance based approach, comparing the VCF files normalized by vt normalize and GATK LeftAlignAndTrimVariants, and comparing the UVCF files produced by UPS-indel. For the distance based approach, two indels are considered equivalent if (1) they belong to the same indel type (either both are insertion type or both are deletion type), (2) have the same base pairs inserted/deleted, and (3) are in close proximity (within ± 5 bps from each other). For the normalized VCF files and UVCF files, the same approach was used as discussed earlier for finding redundant indels.

First the VCF files produced by the five indel calling tools were compared to find overlap among them to determine the number of common indels using the distance based approach. In the second step, the VCF files of the five indel calling tools were normalized using vt normalize and GATK LeftAlignAndTrimVariants separately. For this sample, both normalization tools produced the same normalized VCF files. The normalized VCF files of five indel calling tools were compared to determine the common number of indels. Finally, UPS-indel was used to produce the UVCF files for the five indel calling tools and these UVCF files were compared to determine the common number of indels.

The result shows that the distance based approach found 584 indels in common from the five indel calling tools while 5,514 and 5,575 common indels were found by the normalized VCF and UPS-indel UVCF approaches, respectively. This demonstrates the better suitability of UPS-indel, compared to distance based or existing normalization based approaches, for comparing multiple VCF files. Note that this small number (61) of common indels identified by UPS-indel, but missed by the normalization tools, is based on a single chromosome of a single sample only, and much better performance of UPS-indel would be expected for the whole genome, as observed for the dbSNP and COSMIC datasets.

As mentioned earlier, the tools VarMatch [3], RTG Tools [15], and READDI [16] are also used for comparing indel call sets. However, VarMatch and RTG Tools, which use a branch and bound algorithm, are not suitable for population-scale indel call sets like dbSNP and COSMIC due to densely packed indels in those call sets. READDI processes deletions only. These tools are compared with UPS-indel (using the deletion call set of Platypus containing 14,438 deletions for chromosome 11 of the above mentioned single sample from the 1000 Genomes project as the baseline) on the deletion call sets of Dindel, GATK Unified Genotyper, GATK Haplotype Caller, and Pindel as the query call set to check overlap with the baseline. Table 10 shows the comparison of overlaps.

Table 10 shows that UPS-indel finds more common indels than the state-of-the-art tools when comparing multiple indel call sets. These tools are heuristic and therefore ignore indels that violate a particular heuristic criterion. For example, READDI searches for equivalent indels in an indel's neighboring region defined by the neighborhood size, and RTG Tools uses a cutoff strategy when the search space is too large. UPS-indel, on the other hand, exhaustively searches for and finds all equivalent indels, thus finds more common indels than the aforementioned tools.

## Conclusion

This paper describes UPS-indel, a user friendly tool that creates a universal positioning system called UPS-coordinates for all indels listed in a VCF file, and exhaustively finds all equivalent indels. The UPS-coordinate is a range of positions where all indels equivalent to a specific indel can occur. Since equivalent indels produce the same mutant sequence and thus have the same biological effect, reporting them as separate indels causes data redundancy and may artificially inflate the statistics of indel variations. Under the proposed universal positioning system, all equivalent indels have the same UPS-coordinate which avoids possible annotation ambiguity. Therefore, by checking the UPS-coordinate, one can easily filter out redundant indels from variant databases. UPS-indel is robust enough to handle complex variants and is able to detect more redundant indels than the currently existing approaches. UPS-indel could be widely used for easy and accurate systematic comparison of indels generated by different indel calling programs or deposited in databases. By eliminating the indel redundancy issue, this work offers the community the proposed universal positioning system to represent indels (so as to avoid ambiguity), which can greatly improve various downstream genomic analyses related to indels.

## Additional File

**Additional File 1: Supplementary materials of UPS-indel: a Universal Positioning System for Indels.** This additional file includes an example of redundant indels in dbSNP; an example of equivalent deletion; chromosome wise redundant indel ratio of UPS-indel, vt normalize, BCFtools and GATK LeftAlignAndTrimVariants; chromosome wise Redundant indel ratio of UPS-indel, vt normalize, BCFtools, and GATK LeftAlignAndTrimVariants for COSMIC coding indel dataset; chromosome wise redundant indel ratio of UPS-indel, vt normalize, BCFtools, and GATK LeftAlignAndTrimVariants for COSMIC noncoding indel dataset; a bar graph showing the chromosome wise comparison of redundant indel ratio among UPS-indel, vt normalize, BCFtools, and GATK LeftAlignAndTrimVariants for the dbSNP dataset; a bar graph showing the chromosome wise comparison of redundant indel ratio among UPS-indel, vt normalize, BCFtools, and GATK LeftAlignAndTrimVariants for COSMIC coding indels; a bar graph showing the chromosome wise comparison of redundant indel ratio among UPS-indel, vt normalize, BCFtools, and GATK LeftAlignAndTrimVariants for COSMIC noncoding indels

## Declarations

### Ethics approval and consent to participate

Not applicable

### Consent for publication

Not applicable

### Availability of data and materials

The latest version of dbSNP VCF file can be found here: ftp://ftp.ncbi.nlm.nih.gov/snp/organisms/human_9606/VCF/. VCF file for the COSMIC coding mutation is available at http://grch37-cancer.sanger.ac.uk/cosmic/files?data=/files/grch37/cosmic/v78/CosmicCodingMuts.vcf.gz and non coding mutation dataset is available at http://grch37-cancer.sanger.ac.uk/cosmic/files?data=/files/grch37/cosmic/v78/CosmicNonCodingVariants.vcf.gz. All of these VCF files contain SNPs, Indels, and other types of genetic variants. To extract only indels, we used VCFtools which is available at http://vcftools.sourceforge.net/. The command line version of UPS-indel is available at http://ups-indel.sourceforge.net with the instruction of how to install and use UPS-indel.

## Competing interests

The authors declare that they have no competing interests.

## Funding

This material is based on research sponsored by Air Force Research Laboratory under agreement number FA8650-09-2-3938. The U.S. Government is authorized to reproduce and distribute reprints for Governmental purposes notwithstanding any copyright notation thereon. The views and conclusions contained herein are those of the authors and should not be interpreted as necessarily representing the official policies or endorsements, either expressed or implied, of the Air Force Research Laboratory or the U.S. Government.

## Author contributions

M.S.H. developed the software and conducted the computational experiments. M.S.H, X.W., Z.L. and L.Z. designed and analyzed the experiments. L.W. did the mathematical validation. L.Z. planned and supervised the experimental design. M.S.H, X.W. L.W., and L.Z. wrote the manuscript with input from all authors. All of the authors have read and approved the final manuscript.

## Acknowledgement

The authors thank S. Tithi and V. Vijayan for their comments and suggestions.

